# GeneBreaker - Variant simulation to improve the diagnosis of Mendelian rare genetic diseases

**DOI:** 10.1101/2020.05.29.124495

**Authors:** Phillip A. Richmond, Tamar V. Av-Shalom, Oriol Fornes, Bhavi Modi, Alison M. Elliott, Wyeth W. Wasserman

**Author notes:** These authors contributed equally (Phillip A. Richmond, Tamar V. Av-Shalom). **Corresponding Author** Professor Wyeth W. Wasserman < >.

## Abstract

Mendelian rare genetic diseases affect 5-10% of the population, and with over 5,300 genes responsible for ~7,000 different diseases, they are challenging to diagnose. The use of whole genome sequencing (WGS) has bolstered the diagnosis rate significantly. Effective use of WGS relies upon the ability to identify the disrupted gene responsible for disease phenotypes. This process involves genomic variant calling and prioritization, and is the beneficiary of improvements to sequencing technology, variant calling approaches, and increased capacity to prioritize genomic variants with potential pathogenicity. As analysis pipelines continue to improve, careful testing of their efficacy is paramount. However, real-life cases typically emerge anecdotally, and utilization of clinically sensitive and identifiable data for testing pipeline improvements is regulated and limiting. We identified the need for a gene-based variant simulation framework which can create mock rare disease scenarios, utilizing known pathogenic variants or through the creation of novel gene-disrupting variants. To fill this need, we present GeneBreaker, a tool which creates synthetic rare disease cases with utility for benchmarking variant calling approaches, testing the efficacy of variant prioritization, and as an educational mechanism for training diagnostic practitioners in the expanding field of genomic medicine. GeneBreaker is freely available at http://GeneBreaker.cmmt.ubc.ca.

## Background

Next-generation sequencing, and increasingly third-generation sequencing, has been revolutionary in rare disease diagnosis (Wise et al., 2019). By sequencing the entire genome, millions of variants are identified, prioritized, and then manually curated to arrive at a diagnosis for the affected individuals. This process occurs both in a familial setting (e.g. trio sequencing of mother-father-proband) as well as in groups of individuals with similar phenotypes (e.g. case series or cohort studies). The analysis process can be broken down into two distinct steps: variant calling and variant interpretation.

Advances in bioinformatic tools for variant calling have recently expanded the types of variants being detected from single nucleotide variants (SNVs) and small insertions and deletions (indels), to now routinely include copy number variants (CNVs), tandem repeat expansions (REs), mobile element insertions (MEIs) and complex structural variants (SVs) such as inversions, insertions, and translocations. A major challenge within the expansion to these more complex variant types is the large amount of noise and artefacts stemming from limitations of short-reads—with length of 100-250 base pairs (bp)—attempting to resolve repetitive sequences within the genome. These short reads, even when being sequenced from two ends of a DNA fragment with ~500 bp length, cannot span the repetitive elements in the genome. Specifically, interspersed repeats and short tandem repeats whose length often exceeds 500bp, cannot be resolved and uniquely mapped against a reference. These mapping ambiguities lead to strict region-based filtering in order to mitigate noise, in spite of the significant proportion of disease-associated genes overlapping these regions (Ebbert et al., 2019; Goldfeder et al., 2016). As both read length technology and algorithmic approaches continue to evolve, new tools will emerge as candidates to use within diagnostic pipelines (Wenger et al., 2019). While benchmarking the variant calling process in human genomes has been a focus of international consortia, the majority of comparisons focus on evaluations of healthy individuals with well characterized variant sets (Krusche et al., 2019). In the diagnosis of rare genetic diseases, benchmarks should be focused on assessing the detection capacity of pathogenic or potentially pathogenic variants.

Beyond the landscape of rapidly evolving variant calling approaches is the emergence of a multitude of variant interpretation tools and pipelines. Several *in silico* effect predictors for the assessment of functional variant impact already exist, and research efforts are now adding interpretation capacities for features outside of the coding regions of the genome, e.g. splice regulating sequences (Jaganathan et al., 2019). Furthermore, continuously expanding population databases serve as filters to help identify rare genomic events (Karczewski et al., 2020; Lek et al., 2016). Both *in silico* predictors and the population allele frequency of observed variants are used as filters when attempting to identify a pathogenic variant causal for rare genetic disease phenotypes.

Combinations of different variant calling and variant interpretation pipelines are implemented across the world in clinical-grade and research-grade genomic diagnostic laboratories. These approaches utilize tools which consistently receive upgrades to underlying software and databases used within analysis pipelines. As diagnostic pipelines continue to evolve, there is an emerging need for specific performance testing to compare different tools and ensure each new version of an analysis pipeline can identify the disrupted gene in an applied setting. The process of iteratively “spiking” thousands of pathogenic variants into a background set of variants is a common process, especially within research papers publishing novel prioritization methods (e.g. Exomiser (P. N. Robinson et al., 2014)). However these methods typically draw upon known pathogenic events–usually curated by consortiums such as ClinVar–and are often limited to SNVs and indels. As multiple classes and genic impacts begin to be explored within the automation space, tools which create unique combinations of rare disease scenarios will become necessary for testing. Combining multiple classes of variants across inheritance patterns is critical to rare disease diagnosis, and the careful creation of such scenarios enables benchmarking “edge” cases within automated pipelines.

There is a demand for developing capacity to create synthetic scenarios of rare disease cases for education and training purposes. As genomic medicine advances into standard of care, there will be a need for easy access to training datasets of rare disease cases of increasing complexity. Institutional policies and guidelines around access and use of sensitive and identifiable data for research purposes beyond that of the specific disease diagnosis vary significantly between different studies and globally (Martani, Geneviève, Pauli-Magnus, McLennan, & Elger, 2019; Raza & Hall, 2017). This means, establishing a universal ‘standard’ of bonafide genomes suitable for personnel training and benchmarking is challenging. Further complicating reanalysis of such data is the possibility of uncovering incidental findings, perhaps affecting either the parents or the proband (Green et al., 2013). In light of incidental findings, this can lead to institutional policies which are strict regarding reanalysis of data post-diagnosis. Even with access to such data for educational purposes, the scale or volume of available data would be limited compared to the potentially infinite possibilities of simulated genomic errors.

To meet these needs and serve the growing community of genomic medicine, we developed GeneBreaker: a simulation tool for Mendelian rare genetic diseases. GeneBreaker has an online web interface for designing custom genetic disease scenarios based on user-guided parameters. It has the capacity to simulate variants of multiple different classes, affecting different genic regions, and can either draw upon known pathogenic variants from resources such as ClinVar (Landrum et al., 2014) and ClinGen (Rehm et al., 2015), or facilitate user creation of novel events. Created variants can be embedded within different familial inheritance models, to model real life scenarios which may be encountered within clinical or research settings.

## Implementation

### Architecture

GeneBreaker is a web server deployed as NodeJS, which communicates with a REST API that accesses an underlying data repository (storing gene models and known pathogenic variants), and a variant simulation framework written in the Python programming language. User interaction with the online tool guides the stepwise process of variant simulation as follows: 1) select the gene and transcript to be “broken”; 2) simulate the first variant by selecting variant class and either novel creation or existing pathogenic variant(s); 3) proceed to design the second variant or finish and output a Variant Call Format file (VCF; Figure 1A). Variant creation is done with Mendelian disease cases in mind, where a single gene/locus is disrupted in either a dominant (single variant) or recessive (one variant per allele) manner. Gene models and known pathogenic variants are stored within a MySQL database, which is used for variant creation. Python code interacts with the MySQL database REST API to extract variant and gene information, and subsequently creates variants based on user parameters. Code for variant creation and MySQL database interaction can be found here (https://github.com/wassermanlab/GeneBreaker). While the primary output of the simulator is a VCF file, there is also the capacity to enable downstream benchmarking. Support for downstream benchmarking includes facilitating a transition to a VarSim-compatible VCF file for full synthetic WGS simulation, and integration into background variant sets for testing annotation and prioritization approaches (Figure 1B). Details about the variant simulation process and downstream benchmarking are below.

**Figure 1.**
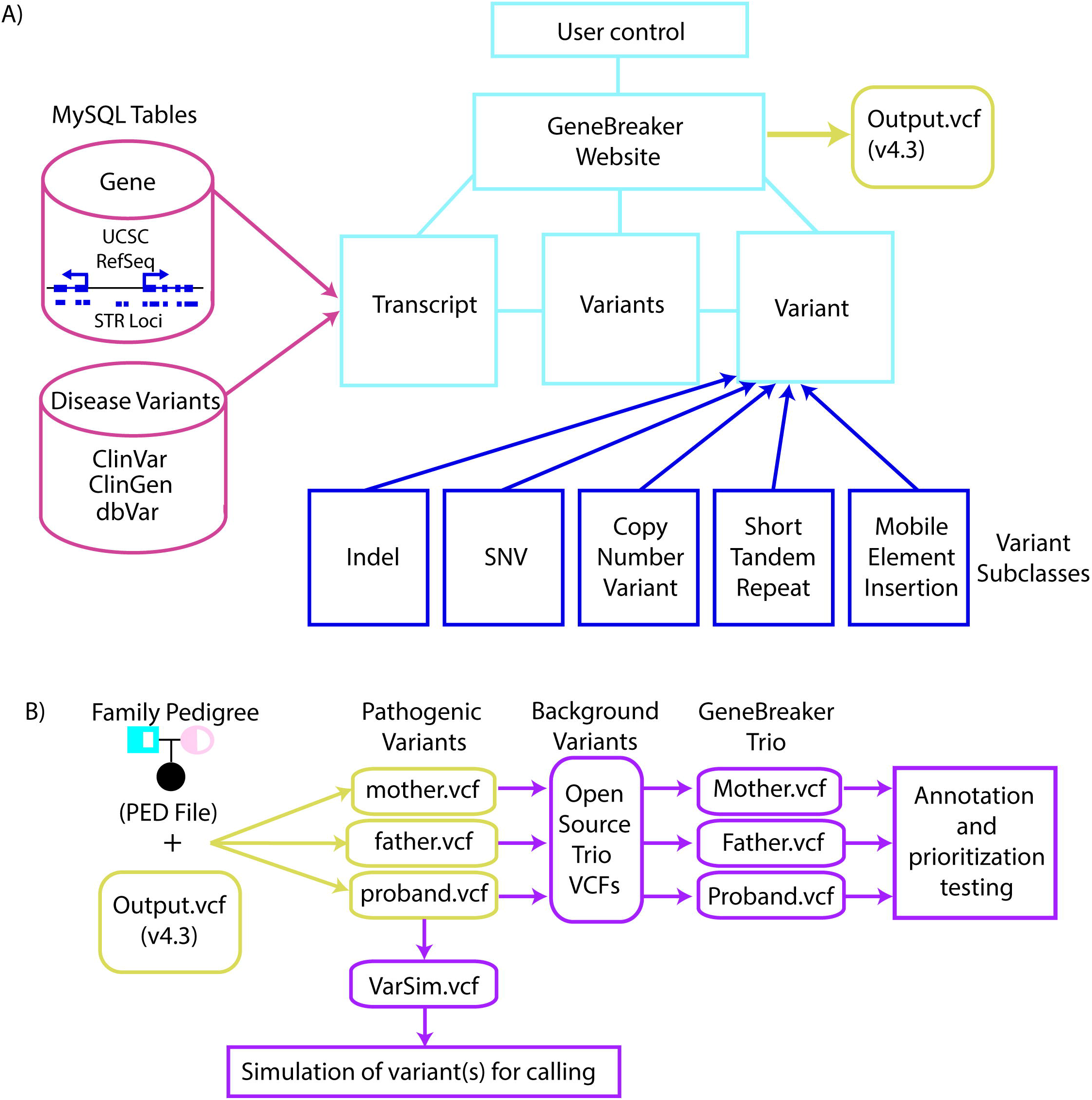
GeneBreaker Overview. (A) Overview of GeneBreaker design framework showing user interaction with the website (light blue), connected MySQL tables (red), underlying variant subclasses (dark blue), and output VCF file (yellow). The user interacts with the GeneBreaker website (light blue) which is connected to hidden components for gene description and variant creation/selection. (B) Downstream benchmarking operations enabled by GeneBreaker including splitting variant amongst VCF files according to user-designed pedigree (yellow), and then either spiking-in the variant within open source trios for annotation and prioritization testing or simulating the proband variant as a full synthetic simulation via VarSim (purple).

### Host web server and underlying data

GeneBreaker is hosted on a virtual webserver at the Centre for Molecular Medicine and Therapeutics, with 12GB of RAM and 4 CPUs running CentOS 7. The Variant data within the underlying repository comes from open source variant catalogues including ClinVar (Landrum et al., 2018), ClinGen (Rehm et al., 2015), and a manually curated set of pathogenic short tandem repeat expansions (Supp. Table S1) (Dolzhenko et al., 2020). Gene models come from the RefSeq annotation database provided by the UCSC Genome Browser (NCBI Homo sapiens Annotation Release 109 (2018-03-29)) (Haeussler et al., 2019).

### Variant simulation walk-through

The user interaction with the simulation process is stepwise and guided, and a video tutorial detailing the creation process is available at: http://genebreaker.cmmt.ubc.ca/more_info. The variant creation process is detailed below.

#### Initial Configuration

Starting at the Variant Designer page (http://genebreaker.cmmt.ubc.ca/variants): 1) User selects reference genome, proband sex, and enters a gene symbol into the ‘gene’ textbox; 2) User clicks “Fetch Transcripts” and all transcripts associated with the gene symbol appear, and the user selects a single transcript which is then displayed in the IGV browser. The primary transcript, as defined by RefSeq Select (https://www.ncbi.nlm.nih.gov/refseq/refseq_select/), appears with an asterisk to guide transcript selection; 3) User proceeds to variant creation by clicking “Next”.

#### Variant Creation

From the Variant 1 Info page: 1) User selects the “Region”, which includes a set of genomic regions: coding, UTR, intronic, and genic (anywhere in the body of the gene). These genomic regions restrict the set of possible variants to those which overlap the defined regions, e.g. coding selection means a variant will have to be created over coding regions of the selected gene; 2) User selects the variant “Type”, either choosing from predefined variant sets: clinvar, clingen copy number variant, and short tandem repeat; or novel creation methods: copy number variant, mobile element insertion, indel, single nucleotide variant; 3) User selects the “Zygosity”: heterozygous or homozygous.

For each of the variant classes, additional information will be required as input. All variant positions are represented in the 1-based half-open coordinate system. Users must choose positions for variants which overlap the selected regions. For *clinvar* and *clingen copy number variant*, clicking the “Fetch variants” box will populate the window with coordinate-sorted variants from the ClinVar or ClinGen databases which overlap the defined genomic region. Clicking on a single variant line will select it and enable the user to proceed to the next page. For *copy number variant*, the user specifies the start and end positions, as well as the copy change (deletion or duplication). For *mobile element insertion*, the user specifies the start position and element type: LINE, ALU, or SVA. For *indel*, the user specifies the start position, and the length as a positive integer for insertion of random nucleotides, or as a negative integer for a deletion. For *single nucleotide variant*, there are multiple SNV types which can be selected: stoploss, missense, nonsense, synonymous, or simply creating alternate alleles from any of the four nucleotides A, T, C, and G. All effects are listed independent of the region selected, however the effects must comply with the region in order to proceed. As an example, selecting intronic region and nonsense variant will give an error. For nonsynonymous variant effects (stoploss, missense, nonsense), the variant position must have the capacity to create an amino acid change. As an example, creating a nonsense (premature stop codon) SNV requires that the altering the single base at the specified position will change the codon sequence to become TAA, TAG, AGA, or AGG. To facilitate this, the three-frame translation in the IGV window can be displayed by zooming into the nucleotide level, and clicking the gear on the right side of the IGV window to select “Three-frame Translate”. For *short tandem repeat*, clicking on the “Fetch Short tandem repeats” box will populate the window with all short tandem repeats which overlap the genic region. Each STR has the repeat motif, and if the repeat is known to be pathogenic then it displays the number of repeat copies which have to be inserted to be considered damaging. After selecting a repeat, enter the repeat length as an integer value of the number of repeats you want to add or remove, using positive or negative integers respectively.

After creating a variant, the user can either repeat the process to create a second variant, or proceed to Family Info to guide the variant inheritance.

#### Inheritance modeling

After the creation of variants, a summary page with Family Info appears and a portion of the variant information is displayed including chromosome, position, reference allele sequence, and alternate allele sequence, adhering to specifications from VCF version 4.2 https://samtools.github.io/hts-specs/VCFv4.2.pdf. Below the variant summary is information for each of the individuals in the family which will be included in the output, starting with information from the proband including sex, presence of variant 1 (Var1), presence of variant 2 (Var2), and affected status (check box). The user will then add family members by clicking on the “Add Family Member” box, choosing to add a mother, father, sister, or brother. After adding the individual, the user will select the boxes for Var1 and Var2 to indicate whether that individual has the variant or not, and their affected status. In the case of homozygous variants, selecting both Var1 and Var2 will add a homozygous variant for that individual. After adding multiple individuals and completing their variant/affected status, the output files can be downloaded by clicking on the “Download Outputs” box. Both the merged VCF containing variant records for all individuals, and the associated pedigree (PED) file can be downloaded.

### Incorporate variants into reads to test detection capacity

Simulation of variants within read sets can be performed in two primary ways: 1) incorporating variants into a reference genome sequence (FASTA format) and then simulating reads from the sequence; or 2) incorporating variants directly into mapped read files. While several tools exist for reference-based incorporation and read simulation, we chose to use VarSim (Mu et al., 2015) due to its ease of use and capacity to simulate other variants alongside the pathogenic variants of interest. Variants were incorporated using VarSim’s default configuration with the hg19 (hs37d5) reference genome, background variants from dbSNP common variants version 150 (https://ftp.ncbi.nih.gov/snp/pre_build152/organisms/human_9606_b150_GRCh37p13/), and DGV variants (from VarSim’s installation script). Before variant incorporation, VCF files from GeneBreaker may need to be reformatted if they contain a mobile element insertion using the reformatForVarSim.py script (https://github.com/wassermanlab/GeneBreaker/blob/master/BenchmarkingTransition/FullSimulation/reformatSimToVarSim.py).

BamSurgeon is a tool for incorporating variants directly into mapped read files (Ewing et al., 2015). GeneBreaker VCF files can be parsed for use within BamSurgeon. We chose to demonstrate the utility of GeneBreaker with VarSim, although both approaches are feasible.

### Spike-in the variant within a trio setting for testing prioritization approaches

Beyond simulating variants for benchmarking variant calling approaches, there is also utility in testing variant prioritization methods. To facilitate this, we collected open source trio PCR-free WGS data from the Polaris project (https://github.com/Illumina/Polaris), and processed it using a standard approach with cutting-edge tools. The data was mapped against the reference genome GRCh37 (http://www.bcgsc.ca/downloads/genomes/9606/hg19/1000genomes/bwa_ind/genome/GRCh37-lite.fa) and GRCh38 (ftp://ftp.ensembl.org/pub/release-96/fasta/homo_sapiens/dna/Homo_sapiens.GRCh38.dna_sm.primary_assembly.fa.gz) using BWA mem (v0.7.17) with default settings (H. Li & Durbin, 2009). Output SAM files were converted into bam and sorted using Samtools (v1.9) (H. Li et al., 2009). Variant calling was done using DeepVariant (0.10.0) (Poplin et al., 2018). Visualizing the mapped reads was done using Integrative Genomics Viewer (IGV) (v2.4.10) (J. T. Robinson et al., 2011). We applied the same mapping and conversion procedure, using the GRCh37 reference genome, to simulated data from VarSim.

The output is a set of VCF files, one per individual, which serve as background variants for both male and female probands (children) and their parents. These variants are deposited in the online repository alongside other full simulations (see Data and Availability). Combining the background variants with the pathogenic variants from the GeneBreaker tool is managed using bash scripts which match the sex and reference genome. These scripts utilize standard tools including bcftools (Heng Li, 2011), htslib (bgzip and tabix), and a custom reformatting script (https://github.com/wassermanlab/GeneBreaker/blob/master/BenchmarkingTransition/BuryVariant/reformatSimToDeepVariant.py). After creating the merged VCF files, we tested them for correct simulation using Exomiser (v12.1.0)(P. N. Robinson et al., 2014). We searched Exomiser output within each gene-based inheritance table for the known gene using the command line tool grep (e.g. “grep -w ABCD1 -n ExomiserOutput_AR_genes.tsv”).

### Creation of training scenarios

Hypothetical cases were created and clinical descriptions generated based on clinical experience. Genes and diseases were selected from OMIM for diverse genetic conditions and include rare disorders, including primary immunodeficiencies, inborn errors of metabolism, developmental disorders, and congenital disorders. HPO terms associated with the disease-associated gene were taken from the HPO website (https://hpo.jax.org/app/), and selected to include common phenotypes associated with the disease.

### Creation of Phenopackets

Phenopackets are an emerging standard, accepted by the Global Alliance for Genomics and Health (GA4GH), for representing phenotypic information in combination with observed variants. For the ten inheritance cases, as well as the ten training scenarios, we created phenopacket JSONs using the phenopacket-schema repository (https://github.com/phenopackets/phenopacket-schema), following the test example for python (https://github.com/phenopackets/phenopacket-schema/tree/master/src/test/python). The Phenopacket JSONs are available on the GeneBreaker website alongside the inheritance testing and training scenarios.

## Results

### Simulator-created rare disease scenarios

We created rare disease scenarios of varying difficulty to test the efficacy of GeneBreaker simulations and for use within benchmarking scenarios (Table 1). These simulations cover different modes of inheritance, variant classes, and genic impacts. The first set of variants covers several inheritance models for combinations of coding variants, either designed by hand or extracted from ClinVar and other published works. Each of the variants in the table were synthetically generated using the online GeneBreaker interface in either the GRCh37 or GRCh38 genome (Table 1). The variants were then assigned to proband, mother, and father according to the inheritance pattern. Lastly, the variants were embedded in the background of open source trios with matched proband sex (Methods). The output from the embedding process is a merged VCF file, which can then be used as input for testing common prioritization workflows, such as the commercial package VarSeq or the open source Exomiser tool (P. N. Robinson et al., 2014).

Beyond creating cases to test inheritance models, we also demonstrate the ability of GeneBreaker to create combinations of variants that are emerging out of anecdotal reports. These variant combinations span multiple classes and genic impacts, and are responsible for the missing heritability in undiagnosed cases (Maroilley & Tarailo-Graovac, 2019). These include a set of cases where pathogenic SNVs and indels lie beyond the coding regions of the gene, and a set of variants which include CNVs, STRs, and MEIs (Table 2).

Lastly, we also designed variants within the “dark regions” of the genome, or regions that are inaccessible to standard variant calling pipelines (Ebbert et al., 2019; Goldfeder et al., 2016). We consider these variants important to simulate due to the need to evaluate results from emerging methods capable of genotyping within such regions (Table 2).

### SNV and indel inheritance testing

To the best of our knowledge, there are no currently available open source tools for prioritizing combinations of different classes of pathogenic variants affecting the same gene. However, the Exomiser tool is a fast and easy-to-use method that can prioritize coding SNVs and indels for Mendelian rare genetic disease cases, and requires as input a merged VCF file, a pedigree (PED) file, and a set of Human Phenotype Ontology (HPO) terms (P. N. Robinson et al., 2008; P. N. Robinson et al., 2014). The HPO terms for a set of selected genes were chosen by matching each gene to a disease using OMIM, and then selecting 4-7 HPO phenotype terms which were common for that disease (https://hpo.jax.org) (Supp Table S2). We simulated each of the inheritance testing cases (Patients 1-10) and searched the output from Exomiser (Methods).

The causal variant was annotated correctly for both user-created SNVs and indels, confirming that our simulation framework for creating novel variants is functional. In seven out of ten cases, the variant was prioritized in the correct inheritance category and was ranked in the top two candidates at the gene level (Table 1). The two scenarios (*SLC6A8* and INPP5E) with compound heterozygous *de novo* inheritance patterns caused issues with Exomiser’s interpretation. A compound heterozygous *de novo* scenario is where a disease-associated allele is inherited from one parent, and a *de novo* mutation disrupts the other allele of the same gene. In both these scenarios, the variants were found to be ranked in the top 10 for dominant inheritance, likely due to the *de novo* variant taking priority. For instance, the *SLC6A8* gene did not show up in the X-linked recessive candidate gene list, but it ranked second in the X-linked dominant gene list. Interestingly, the variant created on the Y chromosome in the *SRY* gene, which is responsible for 46 XY Sex Reversal 1, was not prioritized by Exomiser. It is unclear at which stage this variant was dropped as a result of Exomiser’s inheritance and pathogenicity filtering.

### Testing variant calling

Exomiser is not currently equipped to prioritize CNVs, STRs, and MEIs, and we are not aware of a tool which can integrate these variant classes within inheritance modeling. Therefore, we demonstrate the efficacy of our method by simulating and visualizing a full WGS dataset using the larger variants and the set of variants within the dark regions of the genome (Table 2). Using VarSim, the ten variants from the CNV/MEI/STR and Dark Genome categories were simulated in a single VarSim run. Each of the regions where variants were integrated were visualized with the Integrative Genomics Viewer (IGV) (J. T. Robinson et al., 2011) to validate the variant was simulated correctly at the read level (Figures S1 & S2). An example of the heterozygous duplication of part of the *DMD* gene shows the expected increase in read coverage over the simulated variant (Figure S1A), and the LINE1 transposable element insertion within the intron of *SLC17A5* has the expected signal of soft-clipped reads both upstream and downstream of the insertion site (Figure S1B). For the variants in the dark genome, a homozygous deletion in *SMN2* can be visualized, even though the observed reads are not mapping uniquely to the region (Figure S2A). Lastly, a four base-pair coding deletion within *CFC1* appears in the ambiguously mapped reads, confirming previous reports that specialized methods may be able to locate deletions within these dark and camouflaged regions (Ebbert et al., 2019) (Figure S2B).

### Training scenarios

To emphasize the capacity for GeneBreaker-created scenarios to be utilized within training the next generation of genome medicine practitioners, we created ten hypothetical genetic disorder scenarios. Each hypothetical case has a causal gene, causal variants drawn from ClinVar, HPO terms, and patient descriptions including relevant family history and clinical findings (Table 3, Supp. Mat.). For these ten cases, they can be utilized within educational materials which focus on variant interpretation in rare disease diagnosis.

## Discussion

The diagnosis of rare genetic diseases will continue to improve as novel methods for calling, interpreting, and prioritizing variants emerge and become deployed in a diagnostic setting. Many of the previously challenging variants to genotype, due to variant complexity or existing within “dark” genomic regions, are now regularly identified in WGS datasets. Consequently, there is a critical need for testing the improved pipelines to ensure that such improvements are implemented correctly. While real patient data is paramount for testing the efficacy of a pipeline, access to such data is often prohibitive due to data use restrictions and is limited by the number of observed cases with available data. With simulation, infinite combinations of any possible genomic variant(s) can be designed, from nonsense SNVs, to intronic MEIs, and every combination in between.

The introduction of GeneBreaker addresses an unmet need for simulated rare disease cases in a broadly accessible format. GeneBreaker is deployed as a free-to-use website, enabling user creation of pathogenic variants. Downstream of variant simulation, the tool also supports the transition into benchmarking either variant interpretation or variant calling analysis pipelines. We tested the efficacy of GeneBreaker and the downstream benchmarking transition by simulating rare disease scenarios covering different inheritance models, variant classes, genomic regions, and genic impacts. Using Exomiser, we validated that the variants we simulate have the expected impact, and exposed some limitations in the ability to correctly prioritize variants with challenging inheritance patterns, such as the compound heterozygous *de novo* pattern. Our example using Exomiser highlights that even for combinations of SNVs and indels, inheritance testing can still be improved upon. Larger, more complex variants were visualized in IGV to ensure their correct simulation, confirming that VarSim is a viable option for whole dataset simulation to test variant calling capacity. All of the simulated cases are available online (http://genebreaker.cmmt.ubc.ca/premade_cases), and can serve as a starting point for benchmarking.

GeneBreaker is not intended as another genome sequencing data simulator. Such simulation has been broadly explored over the past decade, with a rich collection of tools available (Escalona, Rocha, & Posada, 2016). The narrow scope of GeneBreaker is placed upon the creation of rare disease simulated genomes, which is achieved by focusing on the generation of diverse forms of genetic disruptions, which can be embedded within real or simulated genome sequencing data. This focus has particular value for two use cases: careful edge-testing of analysis pipelines and training of interpretation specialists.

When it comes to changing software within a clinical diagnostic pipeline—even if it only involves upgrading to a newer version of an existing package—there must be rigorous testing to ensure that the modifications don’t break the pipeline. Any modification to a standard operating procedure must be tested to ensure that it is still capable of performing at or above the existing diagnostic capacity. With the increased adoption of the reference genome version GRCh38, many pipelines currently utilizing older reference genomes will need mechanisms to test their correct functionality in GRCh38 before the transition. Adopting a new reference genome can sometimes have unanticipated side-effects, as was highlighted with an analysis of missing variant calls from realigning WGS datasets in the UK Biobank (Jia, Munson, Lango Allen, Ideker, & Majithia, 2020). A diverse set of variant scenarios involving several inheritance models, genic impacts, and mix of novel and known pathogenic variants has utility for testing these continuously updated pipelines, without the challenge of handling sensitive patient data. GeneBreaker is designed with the goal of creating carefully thought-out edge testing cases, and is not built as a genotype-to-phenotype prioritization benchmarking tool. The process of testing gene-to-phenotype associations and rankings is better handled by considering only known pathogenic events, and utilizing thousands of simulations. The careful creation of complex scenarios is the focus of the benchmarking aspect of GeneBreaker.

GeneBreaker has value beyond benchmarking as a resource for training a new generation of genome analysts. Rare genetic diseases affect a sizable portion of the population, and as WGS moves into the standard of care, many medical professionals will need hands-on training in the utilization of this technology. Having synthetic cases, either at the merged variant set or raw data levels, is imperative. We encourage those developing educational materials for the analysis of rare disease genomes to consider using the simulation capacity of GeneBreaker as a training tool. To emphasize the teaching aspect of GeneBreaker, and to allow rapid adoption into the educational setting, we created ten hypothetical genetic disorder scenarios, complete with phenotypes, patient descriptions, family history, and merged variant sets, all available online (genebreaker.cmmt.ubc.ca). We envision these materials to be invaluable in training individuals working at many institutions, both academic and commercial, who are establishing genome sequencing protocols for rare disease diagnosis.

Future work on GeneBreaker will focus on expanding the simulation capacity to include additional complex variant classes (*e.g.* inversions and translocations) and variants beyond the genic regions known to disrupt gene regulation. Examples of disruptions to regulatory elements include mutations affecting chromatin organization and enhancers (Lupiáñez, Spielmann, & Mundlos, 2016; Smith & Shilatifard, 2014). A major challenge in extending to regulatory elements is that the relevant genomic regions critical for the regulation of a gene are difficult to define. As these genome annotations improve, we look forward to integrating them into GeneBreaker.

We hope that GeneBreaker is adopted by the growing community of researchers and clinicians who are utilizing WGS in the diagnosis of rare genetic diseases. Feedback is appreciated as we continue to improve the software and simulation capacities. GeneBreaker is available for exploration at: http://GeneBreaker.cmmt.ubc.ca.

## Availability of data and materials

Code for the variant simulation, website deployment, and downstream benchmarking can be found on github: https://github.com/wassermanlab/GeneBreaker. The GeneBreaker website can be accessed at: http://genebreaker.cmmt.ubc.ca. Premade variant sets can be found at the GeneBreaker website, and are additionally hosted at Zenodo: https://doi.org/10.5281/zenodo.3829960. Training cases can also be found at the GeneBreaker website, and are additionally hosted at Zenodo.

## Author Contributions

PAR, TVAS, and WWW devised the project. TVAS, and OF coded the database schema for gene and variant extraction. TVAS developed the variant creation software and GeneBreaker website. BM and PAR curated variants for simulation. PAR generated synthetic datasets, analyzed the data, and developed downstream benchmarking capacity. AE developed hypothetical case scenarios. PAR, TVAS and WWW wrote the manuscript, and all authors approved the manuscript.

## Acknowledgements

We would like to acknowledge computational infrastructure support from the Compute Canada and the University of British Columbia Advanced Research Computing organizations. We also thank members of the Wasserman lab for testing the software.

## Web Resources

- NCBI RefSeq Select: https://www.ncbi.nlm.nih.gov/refseq/refseq_select/
- GRCh37 Reference Genome: http://www.bcgsc.ca/downloads/genomes/9606/hg19/1000genomes/bwa_ind/genome/GRCh37-lite.fa
- GRCh38 Reference Genome: ftp://ftp.ensembl.org/pub/release-96/fasta/homo_sapiens/dna/Homo_sapiens.GRCh38.dna_sm.primary_assembly.fa.gz

