## Supplemental Information for "GeneBreaker - Variant simulation to improve the diagnosis of Mendelian rare genetic diseases"

Included in this document are:

- Supplemental Tables
- Supplemental Figures
- Supplementary Patient Descriptions

#### Supplemental Tables

| Variant type | Source | Date Acquired |
|---|---|---|
| STR (Background) | https://github.com/gymreklab/GangSTR | 2019-07-22 |
| STR (Pathogenic) | https://github.com/Phillip-a-richmond/STR_Analysis/tree/master/CompareSTRDatabases | 2019-07-22 |
| Deletion (CNV, Pathogenic) | https://ftp.ncbi.nlm.nih.gov/pub/dbVar/sandbox/sv_datasets/nonredundant/deletions/ | 2019-07-22 |
| Duplication (CNV, Pathogenic) | https://ftp.ncbi.nlm.nih.gov/pub/dbVar/sandbox/sv_datasets/nonredundant/duplications/ | 2019-07-22 |
| SNV/Indel (Pathogenic + Benign + others) | ftp://ftp.ncbi.nlm.nih.gov/pub/clinvar/ | 2019-07-22 |

**Supplemental Table 1**
Data sources for variants within the database available for simulation.

| Inheritance | Gene | HPO terms |
|---|---|---|
| autosomal dominant maternal | JAK1 | HP:0000964; HP:0001047; HP:0001880; HP:0001508; HP:0032064 |
| autosomal dominant de novo | MSH2 | HP:0200008', 'HP:0001250', 'HP:0001276', 'HP:0012378', 'HP:0002027', 'HP:0001276' |
| autosomal recessive homozygous | MALT1 | HP:0000964; HP:0001047; HP:0001581; HP:0004386; HP:0002090; HP:0002205 |
| autosomal recessive compound heterozygous | CFTR | HP:0002613', 'HP:0002721', 'HP:0002024', 'HP:0002206', 'HP:0002205', 'HP:0001738' |
| autosomal recessive compound heterozygous de novo | INPP5E | HP:0001252', 'HP:0001251', 'HP:0001263', 'HP:0002553', 'HP:0001320', 'HP:0002793' |
| X-linked dominant de novo | MECP2 | HP:0001250', 'HP:0001257', 'HP:0005484', 'HP:0002187' |
| X-linked recessive homozygous | WAS | HP:0000964; HP:0001047; HP:0001880; HP:0001508; HP:0032064 |
| X-linked recessive compound heterozygous de novo | SLC6A8 | HP:0001290', 'HP:0001270', 'HP:0000252', 'HP:0000718', 'HP:0008583', 'HP:0000540', 'HP:0008583' |
| X-linked recessive hemizygous de novo | ABCD1 | HP:0001268', 'HP:0001250', 'HP:0000709', 'HP:0008207', 'HP:0002180', 'HP:0002500' |
| Y-linked de novo | SRY | HP:0012245', 'HP:0011969', 'HP:0000032', 'HP:0000098' |

**Supplemental Table 2**
HPO terms for Exomiser testing with the different inheritance patterns, associated to the 10 genes used within the created scenarios.

### Supplemental Figures

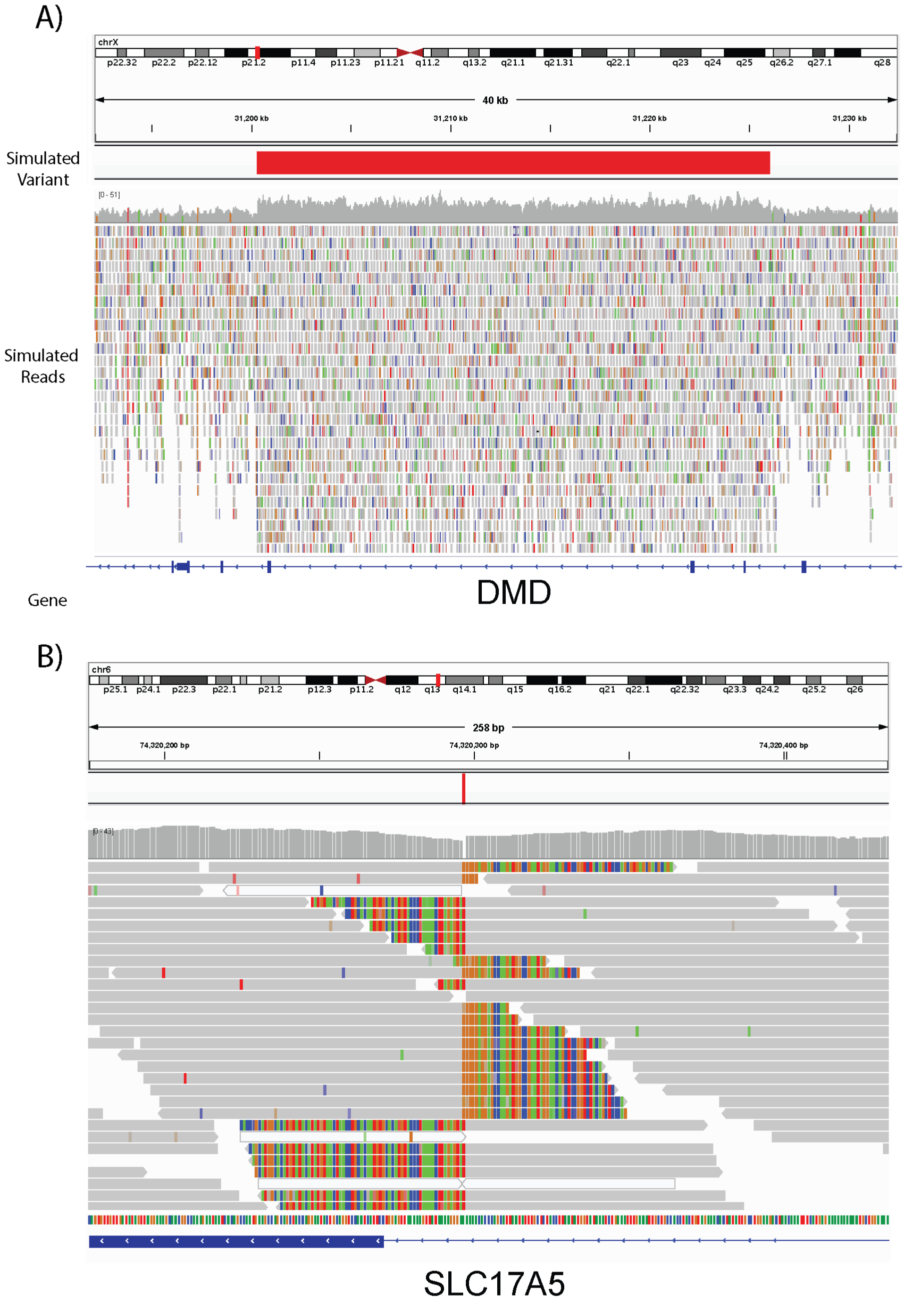


**Supplemental Figure 1**
IGV snapshots of a heterozygous duplication in DMD (A), and a mobile element insertion in SLC17A5 (B). Each image has the simulated variant (red), the mapped simulated reads, and the affected gene.

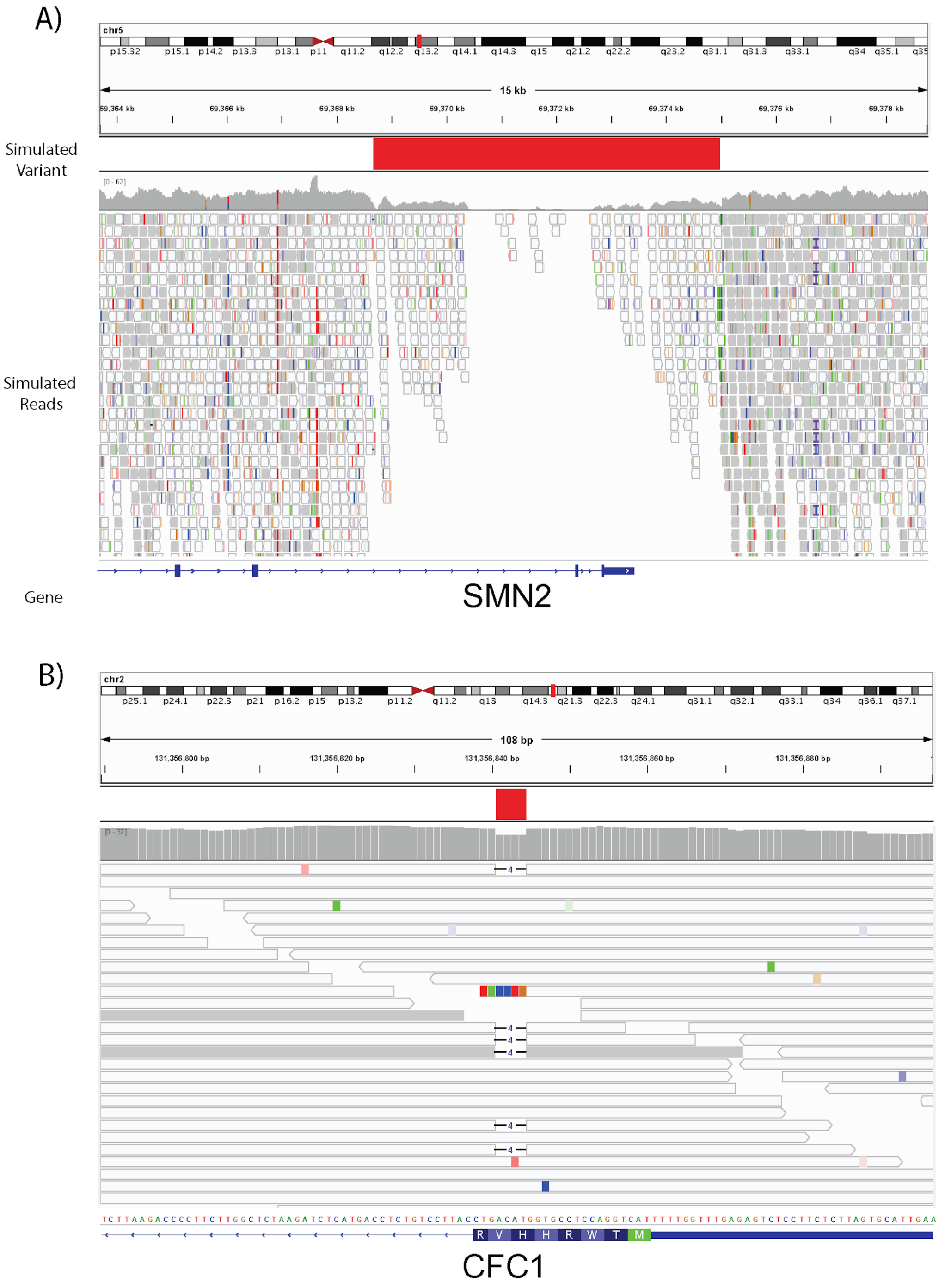


**Supplemental Figure 2**

IGV snapshots of a homozygous deletion in SMN2 (A), and a four base-pair deletion in CFC1 (B). Each image has the simulated variant (red), the mapped simulated reads, and the affected gene.

### Supplementary Patient Descriptions

#### Case 1

##### Clinical and Family history

A two year old boy presents to his pediatrician with bloody diarrhea, eczema, petechiae and easy bruising with no associated trauma. He has had multiple treatments for otitis media (ear infections). Investigations reveal thrombocytopenia (decreased number of platelets) and abnormal morphology (small platelets). The proband is an only child. Both parents do not have any significant medical problems. The family history is significant for a maternal uncle with autoimmune problems and a recent diagnosis of lymphoma.

##### HPO terms & IDs for this patient

Eczema, Thrombocytopenia, Otitis media, Hematemesis
‘HP:0000964', 'HP:0001873',  'HP:0000388', 'HP:0002248'

---

#### Case 2

##### Clinical and Family history

A 41 year old male presents to his family doctor with fatigue and abdominal pain. Initial investigations included a complete blood count, renal and liver panels. He had normocytic anemia and mild elevations of hepatic enzymes. Imaging (CT) of the abdomen demonstrated several hypoattenuating lesions in the liver and edema of the bowel mucosa at the ileocecal junction. Colonoscopy revealed a large mass protruding into the lumen of the distal part of the ascending colon. Subsequent histopathology revealed an adenocarcinoma with negative margins. Microsatellite instability was high. There was no significant family history of cancer or other illnesses.

##### HPO terms & IDs for this patient

Intestinal polyposis, Seizure, Hypertonia, Fatigue, Abdominal pain
HP:0200008', 'HP:0001250', 'HP:0001276', 'HP:0012378', 'HP:0002027'

---

#### Case 3

##### Clinical and Family history

A 14 year old female born to consanguineous parents (first cousins) with short stature and low weight presented with a history of the following: an eczematous rash since (onset when she was a neonate) and persistent severe dermatitis. She has had multiple skin infections and severe inflammatory gastrointestinal disease.

##### HPO terms & IDs for this patient

Eczema, Atopic dermatitis, Recurrent skin infections, Gastrointestinal inflammation, Pneumonia, Recurrent respiratory infections
HP:0000964', 'HP:0001047', 'HP:0001581', 'HP:0004386', 'HP:0002090', 'HP:0002205'

---

#### Case 4

##### Clinical and Family history:

A two year old male presented to his pediatrician with a fever of unknown origin. He was underweight for height and had a history of loose stools. His work up demonstrated an elevated white count. His chest x-ray showed patchy atelectasis (partial collapse of the lung) and a R middle lobe infiltrate. He was admitted to the hospital for IV antibiotics and *pseudomonas aeruginosa* were isolated from culture from a  sputum sample. He underwent sweat chloride testing which was positive. The family history revealed a healthy older sister. Parents were in good health and nonconsanguineous. His mother was of Scottish heritage, and his father was Irish.

##### HPO terms & IDs for this patient

Biliary cirrhosis, Immunodeficiency, Malabsorption, Pulmonary fibrosis, Recurrent respiratory infections, Exocrine pancreatic insufficiency
‘HP:0002613', 'HP:0002721', 'HP:0002024', 'HP:0002206', 'HP:0002205', 'HP:0001738'

---

#### Case 5

##### Clinical and Family history

A two year-old female presented for a well child check. Her early pediatric milestones were reassuring, she walked shortly after her first birthday and began talking around the same time. However, at the appointment, her mother notes that she is no longer using words that she used to and that she no longer feeds herself and seems less engaged. On examination, her growth parameters are in the normal range, but are notable for minimal growth in head circumference over the last 9 months. She verbalizes but does not make intelligible words and does not reciprocate in verbal exchange. She is an only child and the family history is negative with respect to significant medical problems.

##### HPO terms & IDs for this patient

Seizure, Spasticity, Postnatal microcephaly, Intellectual disability profound
‘HP:0001250', 'HP:0001257', 'HP:0005484', 'HP:0002187'

---

#### Case 6

##### Clinical and Family history

A six year old boy is seen by his pediatrician as his mother noted that he has become hyperactive, easily distracted and impulsive. His mother said he was behind with respect to language skills, fine motor skills and coordination. She also expressed concern to the physician that he continues to resist eating solid foods as he frequently chokes and gags when eating. He was a normal appearing male with a slight build. He had increased muscle tone with brisk reflexes at the patellae bilaterally. He was referred to neurology and brain imaging identified symmetric enhancement of signal in the parieto-occipital region with evidence of neuronal loss. The family history was otherwise negative for significant medical concerns. Parents are nonconsanguineous.

##### HPO terms & IDs for this patient

Mental deterioration, Seizure, Psychosis, Primary adrenal insufficiency, Neurodegeneration, Abnormality of the cerebral white matter
'HP:0001268', 'HP:0001250', 'HP:0000709', 'HP:0008207', 'HP:0002180', 'HP:0002500'

---

#### Case 7

##### Clinical and Family history

A seven year old female was referred to Medical Genetics. She had moderate intellectual disability and characteristic craniofacial features that included: arched eyebrows, downslanting palpebral fissures, a prominent nose and micrognathia. She was also noted to have broad thumbs. The family history revealed that her father had significant learning problems (did not finish high school) and similar craniofacial features to his daughter.

##### HPO terms & IDs for this patient

Moderate intellectual disability, arched eyebrows, downslanting palpebral fissures, prominent nose, micrognathia, broad thumbs
‘HP:002342’, ‘HP:0002553’, ‘HP:0000494’, ‘HP:0000448’, ‘HP:0000347’, ‘HP:0011304’

---

#### Case 8

##### Clinical and Family history

A 12-month-old male was seen for routine well child care. His gestation was notable for possible fetal hydrops, but this resolved spontaneously in the beginning of the 3rd trimester. Amniocentesis revealed a normal male karyotype. During the neonatal period, he was identified as having a heart murmur and unilateral cryptorchidism. An echocardiogram was performed and showed a patent ductus arteriosus (PDA) and mild pulmonary valve dysplasia. His developmental milestones were within the normal range, but were at the lower limits of normal for gross motor and language skills. On physical exam he was noted to have short stature with characteristic facial features including low-set, posteriorly rotated ears, widely-spaced eyes, a short and broad nose and wide neck.

##### HPO terms & IDs for this patient

Short stature, congenital heart disease, wide neck, low set posteriorly rotated ears, widely spaced eyes, short and broad nose
'HP:0004322', 'HP:0032318', 'HP:0000475', 'HP:0000368', 'HP:0000316', 'HP:0000445', 'HP:0003196'

---

#### Case 9

##### Clinical and Family history

A ten year old female was referred to Medical Genetics. Her clinical presentation included: Marfanoid habitus, a flat midface, severe myopia and a cleft palate. The family history revealed her father had mitral valve prolapse, sensorineural hearing loss, myopia and a high arched palate. A paternal aunt and the paternal grandfather had hearing loss and near sightedness.

##### HPO terms & IDs for this patient

Marfanoid habitus, flat midface, sensorineural hearing loss, severe myopia, cleft palate, father with mitral valve prolapse, myopia and a high arched palate
'HP:0001519', 'HP:0011800', 'HP:0008625', 'HP:0011003', 'HP:0000175'

---

#### Case 10

##### Clinical and Family history

An 8 year old boy was referred to genetics and had the following clinical presentation: mild intellectual disability, short stature, microcephaly, triangular face, large and prominent ears, hypertelorism, long palpebral fissures and macrodontia. Both parents attended the appointment and the geneticist noted that the mother demonstrated similar craniofacial findings to the son. Upon further questioning and examination, it was revealed that the mother was similarly affected, displaying moderate intellectual disability, and overlap of symptoms with the child.

HPO terms & IDs for this patient

Intellectual disability, short stature, microcephaly, triangular face, large and prominent ears, hypertelorism, long palpebral fissures, macrodontia
‘HP:0001249’, ‘HP:0004322’, ‘HP:0000252’, ‘HP:0000325’, ‘HP:0000400’, ‘HP:0000316’, ‘HP:0000637’, ‘HP:0001572’
